# Single cell RNAseq uncovers a robust transcriptional response to morphine by oligodendrocytes

**DOI:** 10.1101/306944

**Authors:** Denis Avey, Sumithra Sankararaman, Aldrin K. Y. Yim, Ruteja Barve, Robi D. Mitra, Jeffrey Milbrandt

## Abstract

Molecular and behavioral responses to opioids are thought to be primarily mediated by neurons, although there is accumulating evidence that other cell types also play a role in drug addiction. To investigate cell-type-specific opioid responses, we performed single-cell RNA sequencing of the nucleus accumbens of mice following acute morphine treatment. Differential expression analysis uncovered robust morphine-dependent changes in gene expression in oligodendrocytes. We examined the expression of selected genes, including *Cdkn1a* and *Sgk1*, by FISH, confirming their induction by morphine in oligodendrocytes. Further analysis using RNAseq of FACS-purified oligodendrocytes revealed a large cohort of morphine-regulated genes. Importantly, the affected genes are enriched for roles in cellular pathways intimately linked to oligodendrocyte maturation and myelination, including the unfolded protein response. Altogether, our data shed light on a novel, morphine-dependent transcriptional response by oligodendrocytes that may contribute to the myelination defects observed in human opioid addicts.

## INTRODUCTION

Drug addiction is a global health concern associated with considerable adverse impacts. Affected individuals experience severe physical and emotional hardship, which also takes a toll on their families, and society as a whole^1, 2^. The fiscal consequences in the USA alone have been estimated to exceed $600 billion a year (https://www.drugabuse.gov/publications/principles-drug-addiction-treatment-research-based-guide-third-edition). The current epidemic of prescription opioid abuse has contributed to an alarming rise in overdose death rates^2^. Opioid dependent individuals also exhibit increased comorbidity with other psychiatric disorders, including depression, as well as impaired white matter integrity and consequent cognitive deficits^3, 4^. While environmental risk factors certainly contribute to drug addiction, it is well-established that acute drug exposure induces gene expression changes in the central nervous system, and that these changes affect the susceptibility to addiction^5^. All addictive drugs modulate the mesolimbic pathway of the brain by elevating dopamine levels in the nucleus accumbens (NAc), a region in the ventral striatum with a central role in the processing of motivation, reward, aversion and reinforcement learning^6^. Because of this, researchers have aimed to elucidate the effects of a variety of addictive drugs, including methamphetamine^7, 8^, cocaine^8, 9^, and opioids^8–10^, on gene expression in the NAc. However, because the brain is an exceptionally heterogeneous tissue, previous experiments using bulk tissue preparations could not confidently ascribe gene expression changes to a certain cell type, thereby precluding the development of targeted therapies. Recent advances in microfluidics and sequencing technology make it feasible to analyze thousands of single-cell transcriptomes in a single experiment. A variety of single cell RNA sequencing (scRNAseq) platforms have been developed^11–14^, enabling the elucidation of dozens of molecularly distinct central nervous system (CNS) cell types from multiple regions. We profiled the gene expression of 23,276 individual cells from the NAc of mice using Drop-seq^15^ Differential expression analysis uncovered numerous cell-type-specific changes in gene expression due to acute morphine treatment. Unexpectedly, many of the most dramatic morphine-dependent transcriptional responses were observed in oligodendrocytes. We validated these findings by using fluorescence in situ hybridization (FISH) to analyze a subset of highly significant oligodendrocyte- or neuron-specific expression changes. We also performed bulk RNAseq of oligodendrocytes isolated from mock- or morphine-treated mice. Analysis of these data confirmed a remarkably robust transcriptional response to morphine in these cells. Importantly, enrichment analysis of the differentially expressed genes suggests a functional consequence of morphine on oligodendrocyte differentiation and myelination, and provides insights into the underlying molecular mechanisms of opioids. To our knowledge, this is the first study to perform unbiased, cell-type-specific transcriptional profiling *in vivo* following treatment with an addictive drug. Our analyses serve as a framework to study the molecular mechanisms of drugs of abuse using single-cell RNAseq.

### Unbiased single-cell RNAseq analysis reveals 18 distinct cell types in the nucleus accumbens of mice

The molecular responses to morphine and other drugs of abuse by the neurons of the nucleus accumbens (NAc) have been well-studied. However, the responses by specific neuronal or glial subpopulations have not been investigated in an unbiased and high-throughput manner. To address this, we treated mice with morphine or saline as a mock control, and four hours later dissected out the NAc. A single cell suspension was prepared from the dissociated tissue and Drop-seq^15^ was used to obtain transcriptomes from 23,276 single cells (Fig. 1a). We sequenced these cells to an average depth of ~67,000 reads/cell, and detected 1,612 mean genes and 3,228 mean transcripts (Supplementary Fig. 1). Using principal component analysis (PCA) and graph-based clustering (t-SNE)^16^, we identified 18 distinct clusters (Fig. 1b), all of which were present at nearly identical ratios in both mock- and morphine-treated mice, and at comparable ratios among biological replicates (Supplementary Table 1). Next, we assigned a major CNS cell type identity to each cluster (Fig. 1b), by cross-referencing genes that were enriched in specific clusters with published gene expression datasets^17, 18^. Reassuringly, many of the cluster-enriched genes have been previously identified as markers for specific cell types (Supplementary Table 2), and the overall gene expression profiles of our samples correlate well with recent bulk RNAseq^19^ or scRNAseq^17^ analyses of the striatum (Supplementary Fig. 1). Clusters from mock and morphine treated mice exhibited striking similarities in cell number, number of genes detected per cell, and expression levels of cluster-enriched genes (Supplementary Fig. 2), indicating that the overall cell type composition of the NAc is unchanged by acute morphine treatment.

**Figure 1.**
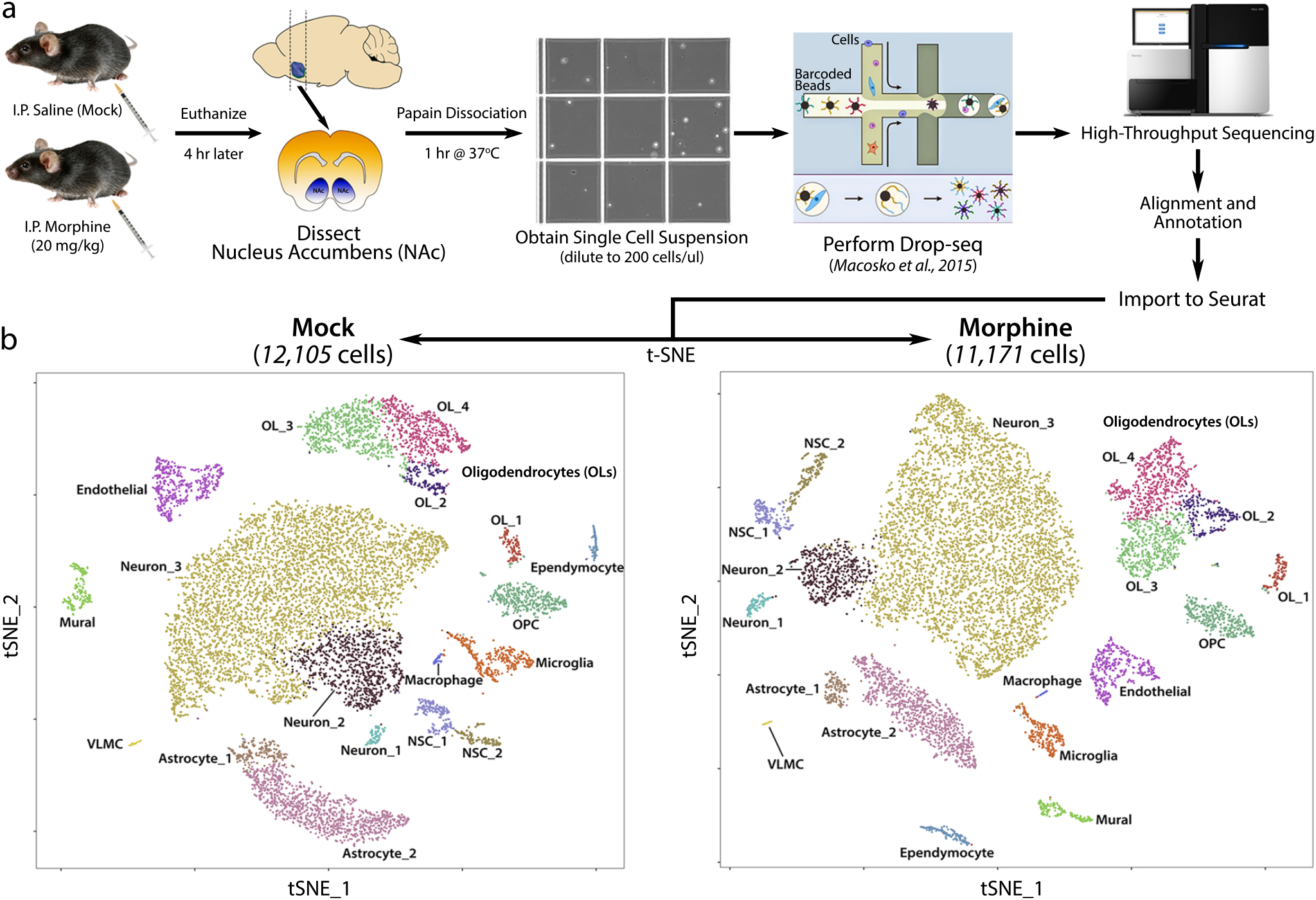
Overview of experimental design and cell types of the mouse nucleus accumbens. (a) Flowchart of experimental design. (b) Spectral t-SNE plots of 12,105 mock or 11,171 morphine cells, colored by density clustering and annotated by cell type identity. OPC: oligodendrocyte progenitor cell; NSC: neural stem cell; VLMC: vascular and leptomeningeal cell.

### Subclustering of neurons enables the identification of a subpopulation of morphine-activated MSNs

The NAc is comprised of many different neuronal subtypes, yet our initial t-SNE analysis resolved only three neuronal clusters. To increase resolution, we subclustered all cells initially identified as neurons, revealing 11 clusters (Fig. 2a; Supplementary Table 3). These clusters directly correspond to the previously identified neuronal subtypes of the NAc. For example, GABAergic D1 and D2 medium spiny neurons (MSNs) are known to comprise ~95% of the neurons in the NAc^20^, and, accordingly, we found two major populations of neurons that were enriched for expression of either *Drd1a, Tac1*, and *Pdyn* (D1 mSns) or *Drd2, Penk*, and *Adora2a* (D2 MSNs) (Supplementary Fig. 3; Supplementary Table 4). Another ~2% of the neurons we identified were GABAergic (*Sst^+^/Npy^+^*) and cholinergic (*Chat^+^/Slc5a7^+^*) interneurons, the latter of which has not previously been characterized by scRNAseq. The ubiquitous expression of *Gad2* (GABA neuron marker) and *Ppp1r1b* (striatal neuron marker) and sparse detection of the glutamate neuron-specific gene *Slc17a7* indicate that our dissections of the NAc were highly accurate, containing minimal contamination with adjacent tissues such as the cortex (Supplementary Fig. 3). Overall, the neuronal subtype diversity that we uncovered is highly concordant with previous studies of cellular diversity in the NAc, including a recently described single-cell analysis of the striatum^17^.

**Figure 2.**
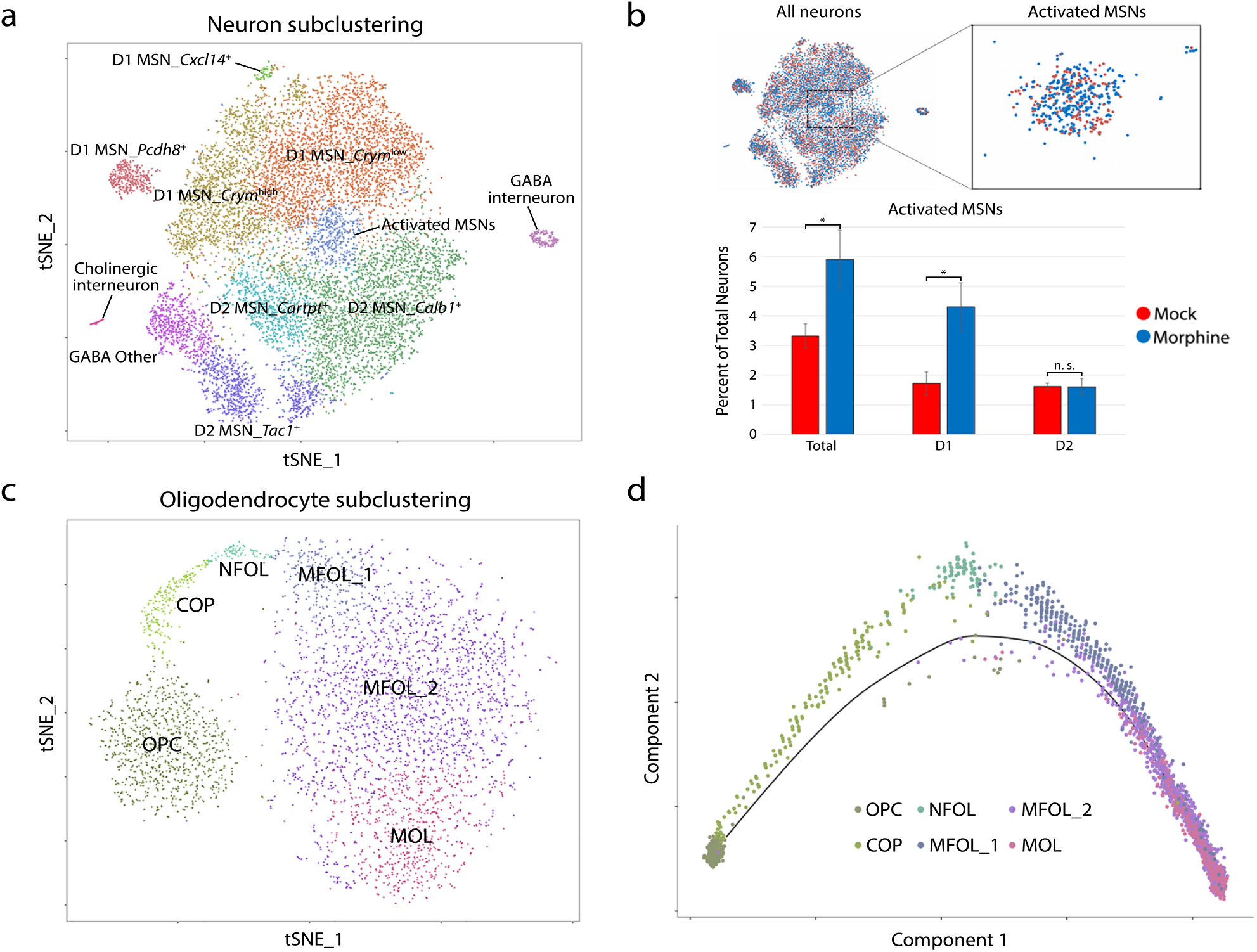
Identification of distinct subpopulations of neurons and oligodendrocytes. (a) Spectral t-SNE plot of 13,033 neurons, colored by density clustering and annotated by cell type identity. (b) Zoomed view of all-neuron t-SNE with cells colored by treatment. Bar graph displays the percent of total neurons in the activated MSN cluster, stratified by treatment and expression of dopamine receptor (D1 vs. D2). Error bars show SD among four biological replicates (*paired t-test, two-sided: p=0.028). (c) Spectral t-SNE plot of 3,817 oligodendrocytes, colored by density clustering and annotated by cell type identity. OPC: oligodendrocyte progenitor cell; COP: differentiation-committed oligodendrocyte progenitor; NFOL: newly formed oligodendrocyte; MFOL: myelin-forming oligodendrocyte; MOL: mature oligodendrocyte. (d) Principal component plot (PC1 vs. PC2) of pseudotime ordering of all oligodendrocytes using Monocle, colored by t-SNE cluster.

We were intrigued by a cluster of neurons in the center of the t-SNE plot, containing both D1 and D2 MSNs expressing high levels of several immediate early genes (Supplementary Fig. 4). Morphine treatment resulted in a statistically significant two-fold increase in the proportion of these “activated” MSNs (Fig. 2b). It has been reported that *c-fos* protein expression is upregulated specifically in D1 MSNs upon acute morphine treatment^21^. Consistent with this, *Drd1a-* expressing cells completely account for the morphine-dependent increase in the proportion of activated MSNs (Fig. 2b). Furthermore, gene set enrichment analysis indicated that genes enriched in this cluster, such as *Fos (c-fos), Junb, Arc*, and *Nr4a1*, are associated with several terms related to opioid addiction, including “Morphine Dependence” and “Opioid-related Disorders” (Supplementary Fig. 5). Thus, we identified a subpopulation of neurons displaying a robust canonical transcriptional response to acute morphine treatment. We performed similar subclustering analyses to further distinguish subpopulations of non-neuronal cells, including astrocytes, immune cells, and vascular cells (Supplementary Table 5; Supplementary Fig. 6). Subclustering of oligodendrocyte (OL) lineage cells resolved them into six clusters, which correspond to developmental states in the continuum from oligodendrocyte progenitor cells (OPCs) to mature OLs (MOLs) (Fig. 2c-d)^22^.

### Acute morphine treatment alters gene expression in a cell-type-specific manner

The primary goal of this study was to identify cell-type-specific alterations in gene expression induced by morphine. To accomplish this, we used SCDE, software that utilizes a Bayesian approach to detect differentially expressed (DE) genes from single-cell data^23^. We presumed that MSNs would exhibit the strongest transcriptional response, since these cells are known to express opioid receptors, display an electrophysiological response to morphine, and mediate morphine-induced behavioral phenotypes. And, indeed, we found that morphine induces the expression of several hundred genes (Supplementary Table 6), including the classical markers *Fos, Junb*, and *Nr4a1*, in D1 neurons of the activated MSN cluster compared to other D1 neurons. Remarkably, we also detected morphine-induced changes in gene expression in nearly every glial cell type (Fig. 3a).

**Figure 3.**
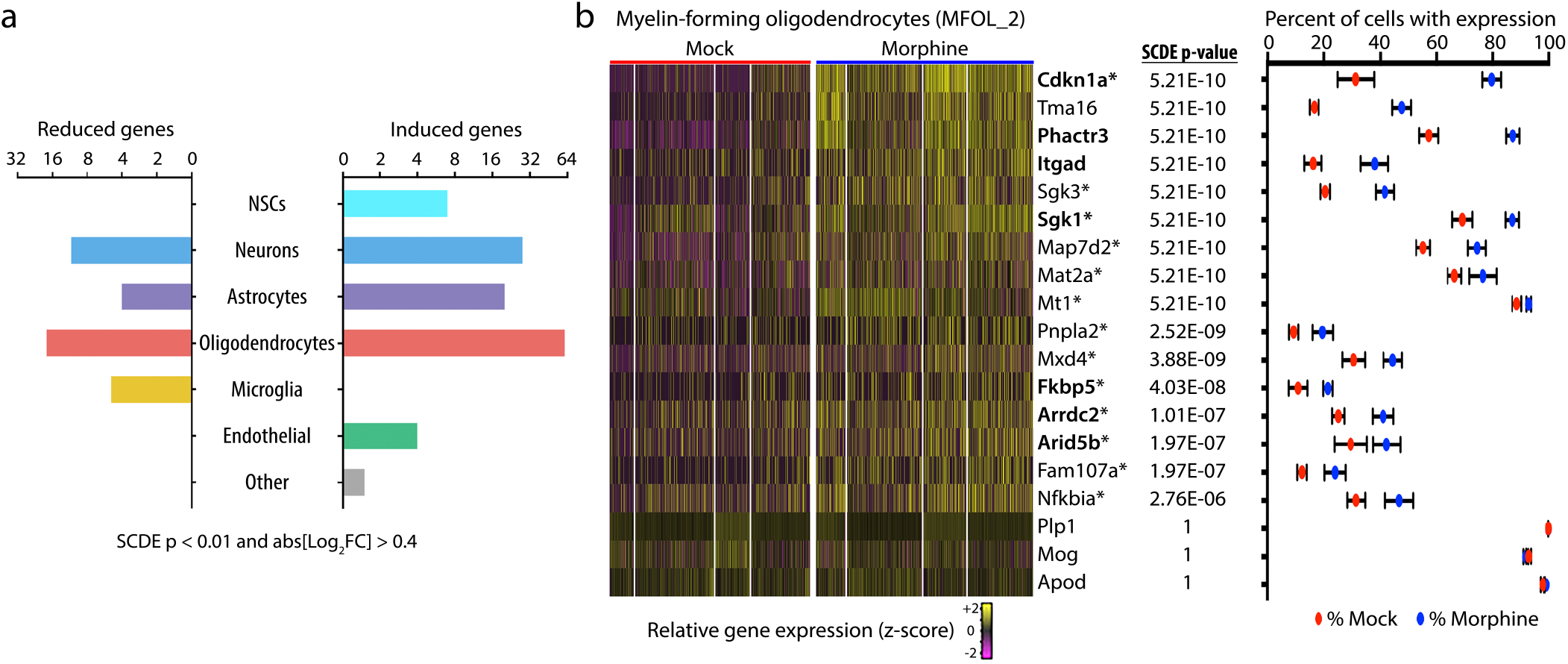
Cell-type-specific differential expression revealed a robust morphine-induced transcriptional response in oligodendrocytes. (a) Numbers of morphine-dependent upldown-regulated genes in each cell type (SCDE p < 0. 01; abs[log_2_FC] > 0.4). (b) Heatmap of relative gene expression for all cells in the largest cluster of myelin-forming oligodendrocytes (MFOL_2) for a subset of morphine-induced genes, ordered horizontally by treatment and biological replicate, and vertically by SCDE p-value. Bolded genes have been previously reported to be induced by morphine, and those marked with an asterisk are reported targets of the glucocorticoid receptor (GR). The right panel shows the percentage of cells in which each gene was detected (mean plus SD of four biological replicates).

Surprisingly, the most numerous significantly DE genes were detected in oligodendrocytes (Fig. 3a), especially myelin-forming OLs (MFOL_2) (Fig. 3b). Many of these genes, including *Cdkn1a, Phactr3*, and *Sgk1*, have been previously reported to be upregulated by morphine in the striatum^8, 10^, but these changes could not be confidently ascribed to a specific cell type (Fig. 3b; bolded genes). Interestingly, 12 of the top 30 morphine-induced genes in OLs (log_2_FC > 0.4; SCDE p-value < 0.0001) are reported targets of the glucocorticoid receptor (GR) (Fig. 3b; marked by asterisk). GR signaling is known to play a key role in morphine-dependent gene expression and behaviors in mice^24^, and more recently has been shown to be critical for a large subset of morphine-induced transcriptional responses in astrocytes^25^. Indeed, we found a significant enrichment of putative GR targets among morphine-induced genes in astrocytes, but also uncovered a distinct subset of such genes in OLs (Supplementary Fig. 7; Supplementary Table 7). Together, these data suggest a robust and previously underappreciated transcriptional response to morphine in oligodendrocytes.

### In vivo validation of cell-type-specific and morphine-dependent gene expression

Although single-cell mRNA profiling is widely used for the identification of new cellular subtypes, calling differentially expressed genes from these data can be challenging due to technical issues such as dropout and dynamic range compression^26^. Therefore, we sought to independently validate some of the most dramatic morphine-dependent changes in gene expression. We used fluorescence *in situ* hybridization (FISH) to co-stain *Cdkn1a* or *Sgk1* and *Mbp* (myelin basic protein), a highly expressed marker of OLs (Fig. 4a). *Cdkn1a* and *Sgk1* signal was sparsely detected in mock-treated mice, while morphine treatment elicited a dramatic increase in puncta number and area that was especially apparent in oligodendrocyte-dense regions, such as the anterior commissure (Fig. 4b) and corpus callosum (Supplementary Fig. 8). This suggests that the OL response to morphine is likely not restricted to the NAc. When mice were pre-treated with naltrexone (NTX), an opioid receptor antagonist, the morphine-dependent induction of *Cdkn1a* and *Sgk1* was dramatically reduced (Fig. 4b), indicating that their induction occurs in an opioid receptor-dependent manner. As expected, morphine-dependent induction of the immediate-early genes *Fos, Junb*, and *Nr4a1* was observed in the NAc and dorsal striatum, predominantly in *Drd1*-expressing cells (D1 MSNs), and was similarly reduced by NTX pre-treatment (Supplementary Fig. 9). Together, these results validate the cell-type-specific transcriptional responses we identified by single-cell RNA-seq, and suggest that both neuron- and OL-specific changes are mediated by opioid receptor activation.

**Figure 4.**
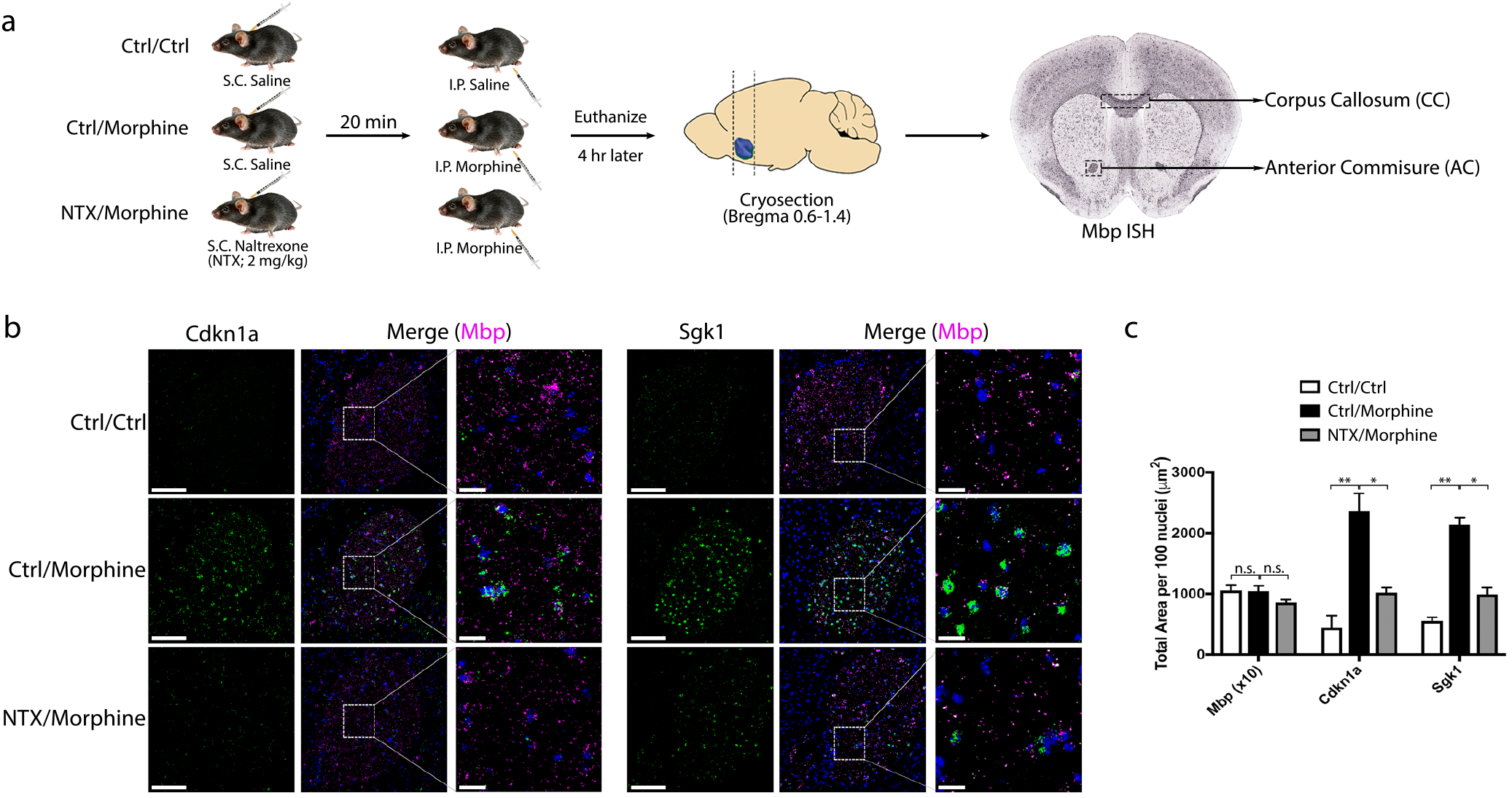
In vivo validation of oligodendrocyte-specific morphine-dependent changes in gene expression by FISH. (a) Schematic of experimental design. Pre-treatment with naltrexone (NTX) should block molecular/behavioral responses to morphine. (b) Representative images of the anterior commissure. Signals are pseudocolored by expression of *Cdkn1a/Sgk1* (green) or *Mbp* (magenta). Scale bars, 100 μm (left panels); 20 μm (right panels). (c) Puncta area quantification for *Mbp* (divided by 10), *Cdkn1a*, and *Sgk1*. Values are shown as mean plus SD (n = 3 mice per condition. ANOVA: n.s. - not significant; **p<0.001; *p<0.01).

### Further characterization of the transcriptional response to morphine by bulk RNAseq of FACS-isolated oligodendrocytes

Because our Drop-seq and FISH data indicated a robust oligodendrocyte transcriptional response to morphine *in vivo*, we sought to further characterize this phenomenon. We adapted an immunostaininglFACS-based approach^27^ to isolate OLs from the entire brains of six mice, three each mock- or morphine-treated. Salinelmorphine injections and tissue dissociation were performed exactly as in our single-cell RNAseq experiments to enable cross-validation between the two datasets. We gated out CD45^+^ and PDGFRa^+^ populations to remove immune cells and OPCs, respectively, and then sorted for myelinating OLs based on their dual expression of GALC and MOG (Supplementary Fig. 10). RNA was purified from each of the six samples and examined by qRT-PCR. We found a 10-20-fold enrichment of the OL markers *Cnp* and *Mog*, and a corresponding de-enrichment of the neuronal marker *Rbfox3/NeuN* and the astrocytic marker *Gfap* in these samples, illustrating the high purity of OLs isolated using this approach (Supplementary Fig. 11). Assured that this was an enriched OL population, we tested the morphine-dependent upregulation of *Cdkn1a* and *Sgk1* by qRT-PCR and found that they were induced by more than 10-fold. (Supplementary Fig. 11).

To ascertain the full extent of morphine-dependent transcriptional regulation in OLs, we next performed RNAseq on these sorted oligodendrocyte samples and compared these results to our single-cell data. We reasoned that the FACS-sorted cell population should be most comparable to the three mature OL clusters observed in our single-cell transcriptional profiling (MFOL_1, MFOL_2, and MOL; collectively referred to as “MO_all”), since these clusters all specifically and ubiquitously express *Mog*, one of the extracellular markers we used to sort the OLs. Strikingly, differential expression analysis of the bulk FACS-RNAseq data confirmed 85.5% (59/69) of the changes detected by SCDE of oligodendrocytes (MO_all). In addition, we uncovered an additional 935 genes significantly affected by acute morphine treatment (DESeq p-adj < 0.05 and abs[log_2_FC] > 0.5; Fig. 5a; Supplementary Table 9). These results demonstrate that the vast majority of morphine-dependent expression changes detected by single-cell RNAseq were indeed *bona fide*; additionally, our bulk RNAseq experiments identified a large set of expression changes that were not observed in our single-cell experiments, illustrating the increased sensitivity of the FACS-RNAseq approach. Strikingly, genes affected in oligodendrocytes comprise nearly a third of all morphine-regulated genes identified by a previous whole-genome microarray analysis of the striatum (Fig. 5b)^8^. This underscores the utility of cell-type-specific approaches for molecular profiling of the brain.

**Figure 5.**
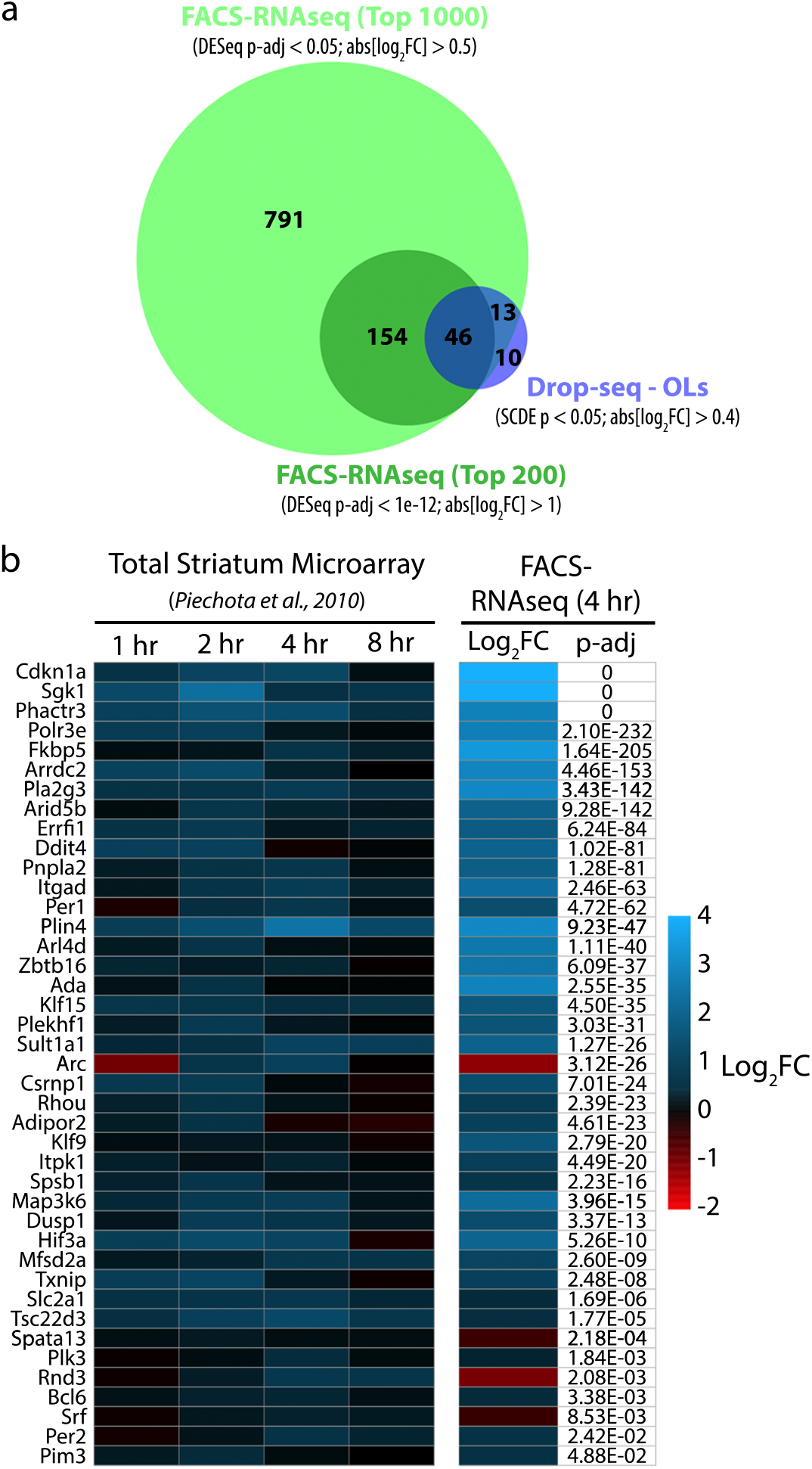
Bulk RNAseq of FACS-isolated oligodendrocytes reveals a remarkable extent of transcriptional regulation by morphine. (a) Venn Diagram illustrating significant overlap between DE genes identified from Drop-seq (MO_all) or FACS-RNAseq. (b) The left panel of the heatmap shows the relative expression (log_2_FC morphine vs. saline at the indicated time point) of a subset of genes previously reported to be induced by morphine, as assess by microarray of total striatum^8^. All of these genes are significantly affected by morphine in oligodendrocytes, according to our FACS-RNAseq analysis.

### Pathway analysis of oligodendrocyte-specific morphine-modulated genes

To obtain biological insights into the response to morphine by OLs, we performed gene set enrichment and pathway analyses. This confirmed the enrichment of GR targets, including all of the GR-related genes we identified by Drop-seq, among the morphine-induced genes in OLs (Supplementary Fig. 12; Supplementary Table 10). We also found that several morphine-repressed genes encode endoplasmic reticulum (ER) chaperone proteins with critical functions in ER quality control (ERQC) and the unfolded protein response (UPR; Fig. 6). Myelin-forming OLs are thought to be particularly susceptible to ER stress, due to their high rates of protein and lipid synthesis^28^. Accordingly, components of the UPR machinery are dysregulated in a variety of demyelinating diseases of the peripheral and central nervous system^28^. The morphine-dependent downregulation of several key components of ERQC/UPR would likely lead to an increase in the intracellular accumulation and/or export of misfolded proteins, which may influence oligodendrocyte survival or myelin composition. The possibility of a morphine-dependent modulation of oligodendrocyte myelination has considerable clinical implications in light of several recent studies describing impaired white matter integrity in the brains of human opioid users^3,4,29–31^.

**Figure 6.**
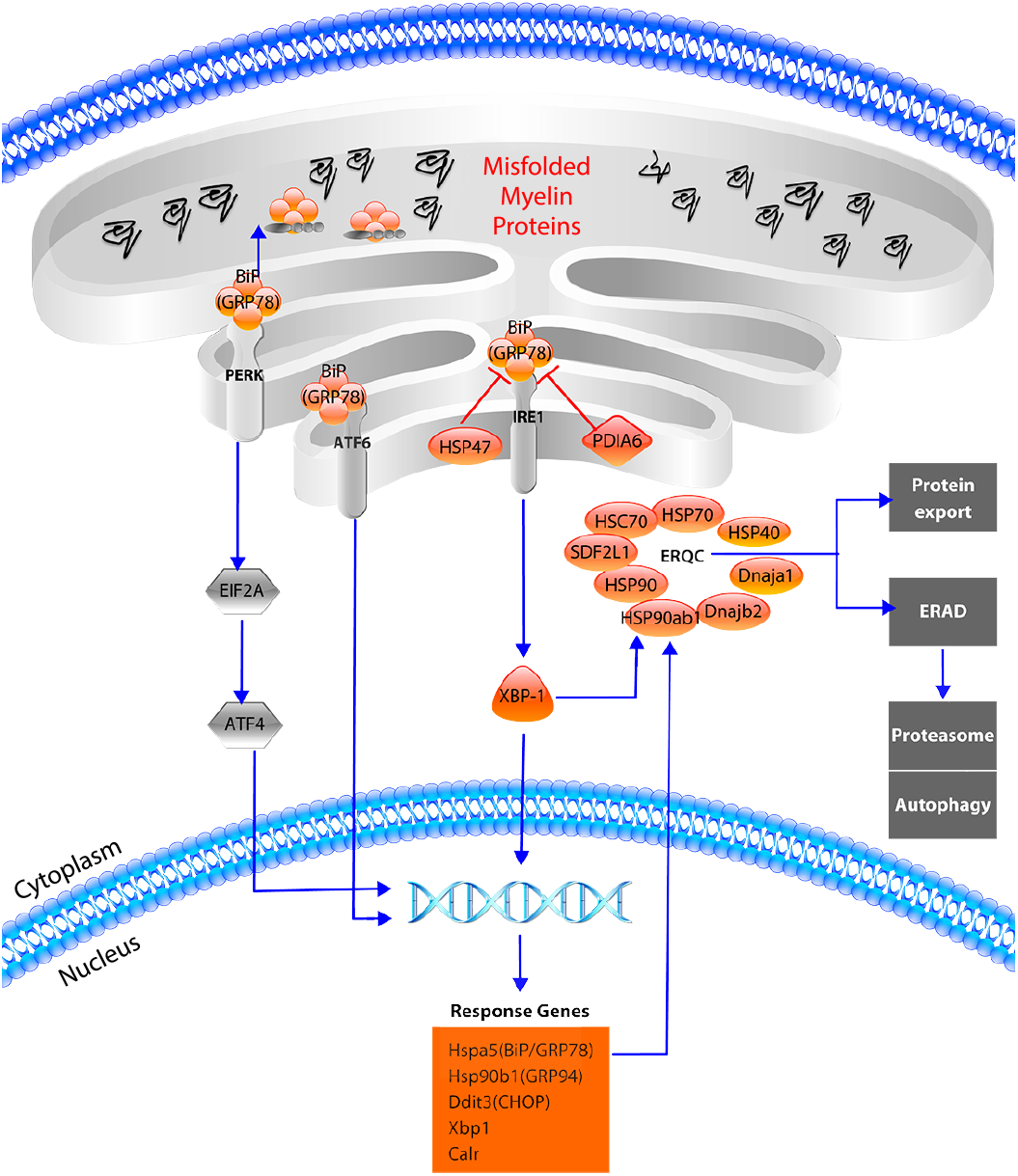
Several components of the unfolded protein response (UPR) are downregulated by morphine in oligodendrocytes. Genes downregulated by morphine in OL based on FACS-RNAseq data are shown in orange while unaffected factors are shown in grey. Notably, the IRE1-XBP-1 axis of the UPR appears to be primarily affected. XBP1, as well as numerous XBP-1-induced genes (shown in orange box) are reduced, as are several endoplasmic reticulum (ER) chaperone proteins involved in ER quality control (ERQC). Under normal conditions, these chaperones will assist in the proper folding and export of proteins, or target misfolded proteins to the proteasome by a process known as ER-associated degradation (ERAD). The downregulation of these factors might lead to the increased accumulation and/or export of misfolded proteins, including myelin constituents. This diagram was generated using ePath3D.

## DISCUSSION

Although most studies to date have focused on the neuronal response to morphine, there is ample evidence to suggest oligodendrocytes also play an important role. For example, it has long been known that mouse oligodendrocytes express both mu- and kappa-opioid receptors (MORs and KORs) in a developmentally regulated sequence^32, 33^. Interestingly, perinatal exposure of rats to buprenorphine or methadone, two currently approved opioid medications, has been shown to regulate OL development and myelination^34, 35^. Recent clinical evidence also strongly suggests dysregulation of OLs in the context of opioid abuse. In particular, brain imaging of heroin users has revealed impaired white matter integrity indicative of myelin pathology^3, 4, 29–31^. Heroin injection has also been linked to a rare form of leukoencephalopathy^53^. Our data indicate that even a single dose of morphine alters the expression of genes in OLs with known functions in differentiation and myelination. We expect that some of these changes would persist upon long-term opioid treatment, and thus our results provide a plausible mechanism for the impairment of OL function with chronic morphine use.

Morphine is known to rapidly increase the serum concentration of corticosterone^36^,which in turn has been shown to induce expression of *Sgk1* (serum/glucocorticoid-regulated kinase 1)^37^. Glucocorticoid receptor (GR) signaling is critical for morphine-dependent gene expression and behaviors in mice^24^, but the specific contributions of OLs have not been examined. Presumably, the morphine-dependent induction of *Sgk1* and a subset of the other genes we identified occurs via transcriptional activation by the GR. This is supported by the significant enrichment of GR target genes, including *Sgk1, Sgk3, Fkbp5, Zbtb16*, and *Nfkbia*, among DE genes identified by both single-cell and FACS-RNAseq analyses of OLs. The robust induction of *Sgk1* we observed is particularly intriguing, since this gene has been recently shown to facilitate OL morphological changes that are associated with depressive-like behaviors^38^. Thus, the GR-*Sgk1* signaling axis in OLs may underlie the comorbidity observed between opioid addiction and depression. It was previously suggested that morphine- and GR-dependent gene expression changes (including *Sgk1* induction) are confined to astrocytes^25^. Our single-cell analyses indicate instead that some genes are induced uniquely in either astrocytes (i.e. *Tsc22d3, Gjb6*) or OLs (i.e. *Sgk1, Sgk3*), while others are induced in both of these cell types (i.e. *Fkbp5, Nfkbia*).

The most significantly downregulated pathway is comprised of genes encoding heat-shock proteins, ER chaperones and other factors critically involved in the unfolded protein response (UPR) and ER quality control (ERQC). Notably, the expression of heat-shock proteins, including Hsp70 (*Hspa1a*) and Hsc70 (*Hspa8*), has previously been shown to be modulated by morphine treatment of rats^39, 40^. Proper regulation of the UPR is essential for efficient myelination, as evidenced by the apparent dysregulation of UPR in multiple myelin disorders, including Charcot-Marie-Tooth disease, Vanishing White Matter Disease, and multiple sclerosis^28, 41^. Under normal conditions, GRP78lBiP (encoded by *Hspa5*) is thought to initiate the UPR by sensing misfolded proteins in the ER^41^, and its conditional ablation in oligodendrocytes or Schwann cells results in cell death and severe hypomyelination^42^. Coincidentally, *Hspa5* is also the most significantly morphine-repressed gene that we detected in OLs. Interestingly, the reduced expression of downstream response genes, including the antisurvival gene CHOP (*Ddit3*) and several ER chaperone proteins, suggests that morphine potently suppresses the canonical UPR, which normally entails attenuation of global protein synthesis, degradation of misfolded proteins, and, under severe ER stress, autophagy or apoptosis. Based on our data and previous studies^34, 35, 43^, opioids appear to promote myelination, which seems counterintuitive to the myelin defects observed in human opioid addicts. Perhaps in the context of chronic opioid exposure, persistent upregulation of myelin synthesis and concomitant downregulation of ERQC leads to increased export of misfolded myelin constituents and therefore altered myelin composition. The integrity and composition of myelin have not been investigated in models of long-term opioid treatment.

In summary, we used single-cell RNAseq to gain insight into cell-type specific changes in the nucleus accumbens in response to morphine. We identified a novel response by oligodendrocytes, and shed light on the underlying molecular mechanisms of this response, specifically implicating GR signaling and the UPR as potential links between the observed gene expression changes and opioid-related disorders. Our studies highlight the utility of single-cell RNAseq for cell-type-specific DE analysis, but also demonstrate the importance of supplementing such data with orthogonal approaches, such as FISH and bulk RNAseq. Our single-cell and bulk RNA-seq datasets should serve as a valuable resource for future research aiming to assess transcriptional responses to opioids (or other drugs of abuse), perform cell-type-specific molecular profiling, or optimize single-cell DE analysis. Future studies are necessary to further investigate the mechanisms, functional consequences, and physiological significance of oligodendrocyte-opioid interactions.

## METHODS

### Mice

All animal care and experimental procedures were approved in advance by the National Institute of Health and Washington University School of Medicine Institutional Animal Care and Use Committee (Protocol #20170030). Male C57Bll6 mice (6-12 weeks old) were used for all experiments, and all animals were socially housed on a 12 h light dark cycle with water and food *ad libitum*. Mice were injected with saline or morphine sulfate (20 mglkg) intraperitoneally, andlor naltrexone hydrochloride (2 mglkg) subcutaneously.

### Dissection and tissue dissociation

Four h after saline (mock) or morphine injection, mice were anesthetized by isoflurane and euthanized by decapitation. Brains were rapidly extracted, cooled in ice-cold Hibernate A (-Ca) media for 5 min, then placed ventral surface up onto a chilled petri dish on ice. The rostral and caudal boundaries of the nucleus accumbens were visually approximated and a 1.5 mm-thick coronal section was obtained using a chilled razor blade. The nucleus accumbens was microdissected at its dorsolateral borders and pooled by experimental condition (two mice per replicate). Pooled tissue samples were digested in papainlDNase I solution for 1 h at 37 °C, according to a previously published protocol^44^. The final volume was 1 mL per mouse nucleus accumbens. Digestion was stopped by dilution with EBSS #2^44^, which contains ovomucoid protease inhibitor. Tissue was centrifuged at 300g for 5 min, resuspended in EBSS #2, then gently triturated with a P200 pipette. The resulting cell suspension was divided between two microfuge tubes and diluted with Drop-seq buffer to a volume of 1 mL per tube (per 2 pooled samples). Drop-seq buffer consisted of 5% (wt/vol) trehalose, Hank’s buffered salt solution (HBSS; magnesium- and calcium-free), 2.13 mM MgCl_2_, 2 mM MgSO_4_, 1.26 mM CaCl_2_, 1 mM glucose and 0.02% bovine serum albumin. Cells were washed twice by centrifugation at 300*g* for 5 min, followed by resuspension in 1 ml Drop-seq buffer. Cells were then passed through a 40-μm mesh filter, diluted to 210 cells/μL using cell concentration estimates from a hemocytometer, and kept on ice until use (<30 min).

### Drop-seq library generation, read alignment and generation of digital expression data

Drop-seq was performed as previously described^45^. Mouse nucleus accumbens suspensions were processed through Drop-Seq in eight separate groups over four separate batches (one batch per day). Libraries were sequenced on the Illumina HiSeq3000 or HiSeq2500. Read 1 was 25 bp (bases 1-12 cell barcode, bases 13-20 UMI - unique molecular identifier), read 2 (paired end) was 75 bp, and the index primer was 7 bp. Raw sequence data was first filtered to remove all read pairs with a barcode base quality of less than 10. The second read (75 bp) was then trimmed at the 5’ end to remove any TSO adaptor sequence and at the 3’ end to remove poly(A) tails of length 6 or greater, then aligned to the mouse (mm10) genome using STAR v2.4.2a with default settings. Uniquely mapped reads were grouped by cell barcode. A list of UMIs in each gene, within each cell, was assembled, and UMIs within ED = 1 were merged together. The total number of unique UMI sequences was counted, and this number was reported as the number of transcripts of that gene for a given cell.

All cell barcodes in which 500 or more genes *and* 1000 or more transcripts (UMIs) were detected were used in downstream analysis. Raw digital expression matrices were generated separately, then the replicates from each experimental group were merged together into a single matrix. Cells with high expression levels of one or more marker gene(s) of more than one cell type (i.e. neuron, oligodendrocyte, astrocyte, endothelial cell, etc.), representing likely cell doublets/multiplets, were removed (~4% of all STAMPS), resulting in a 12,105-cell barcode data set for mock treatment and a 11,171-cell barcode data set for morphine treatment. Gene expression was normalized in Scater^46^ using the method described by Lun et al.^47^, and genes with expression in ≥50 cells were retained for clustering. Before clustering, we removed unwanted variation due to number of UMIs, ratio of reads mapping to mitochondrial genes, and batch effects using Seurat software^48^.

### Unsupervised dimensionality reduction and clustering

We used Seurat to perform clustering^48, 49^. We identified genes that were most variable across the entire data set, controlling for the known relationship between mean expression and variance. We calculated the mean and dispersion (variance/mean) for each gene across all cells, then identified outlier genes. To distinguish principal components (PCs) for further analysis, we plotted the cumulative variance accounted for by each successive PC. Using the PCElbowPlot() function, we found that PCs beyond 15 accounted for negligible variance. We identified molecularly distinct clusters using the FindClusters() function with the Louvain algorithm with multilevel refinement, using a high resolution (2.0) to detect weakly distinguishable clusters. We then built a classification hierarchy, merged clusters with higher than 0.05 Out-of-bag error (OOBE) from a random forest classifier, and performed *t*-distributed stochastic neighbor embedding (t-SNE)^16^ in Seurat with default settings. This allowed us to assign cells into a total of 18 cell type clusters each for mock and morphine. For subclustering, cells corresponding to neuronal, oligodendrocyte, or other clusters were combined, then iteratively clustered as above. Altogether, we identified 29 molecular distinct cell types, all of which were shared between mock and morphine treated mice.

### Single-cell differential expression analysis and pseudotime analysis

To analyze differential gene expression between morphine and mock datasets, we performed pairwise differential expression analysis on specific cell subpopulations using the SCDE package, which fits individual error models to account for stochastic detection of lowly expressed genes^23^. We included all Seurat-generated clusters containing >10 cells from each of 8 biological replicates (Mock1-4, Morphine1-4), and that had relatively uniform distribution across all replicates. Only 3 neuronal subtypes (cholinergic interneurons, D1 MSN_*Cxc/14^+^*, and D1 MSN_*Pcdh8^+^*) did not meet these criteria. We used default settings with the exception that we varied the min.detected parameter of the clean.counts function to filter out genes detected in less than 50 total cells. We included the results of every cluster for all gene with p-value < 0.5 in Supplementary Table 6. For pseudotime analysis, we considered only the top 2000 most highly variable genes (between oligodendrocyte subclusters). We ordered all oligodendrocytes in pseudotime using Monocle with default parameters, except that variation due to biological replicates was regressed out using the residualModelFormulaStr parameter in the reduceDimension function. Cells were colored by their Seurat t-SNE cluster identity.

### In situ hybridization

Morphine and mock treatment was performed as described above, except that mock treatment was preceded by a subcutaneous (s.c.) saline injection (Ctrl/Ctrl), and morphine treatment was preceded by injection with s.c. saline (Ctrl/Morphine) or naltrexone (NTX/Morphine). After 4 h, brains were harvested and immediately flash-frozen in OCT (optimal cutting temperature compound) in liquid nitrogen. We stored frozen tissue blocks at -80°C prior to sectioning. Twelve-micron thick coronal sections were cut at -20°C and adhered to positively charged Leica slides (one of each of the three treatment conditions per slide), then stored at -80°C. Frozen tissue sections were washed briefly with 1X PBS and fixed in 3.7% para-formaldehyde for 1 h. Following fixation, slides were washed twice with 1X PBS and then submerged in 70% ethanol overnight at 4°C for permeabilization. RNA FISH was performed according to the Affymetrix ViewRNA protocol.

### Imaging and image analysis

We imaged FISH AND IHC slides on a Zeiss LSM 880 II confocal microscope at the Washington University Center for Cellular Imaging (WUCCI). Image files were processed and analyzed in Fiji/ImageJ^50^ software. RGB channels corresponding to the two Affymetrix probes and DAPI were split. RNA particles in each channel were then identified in a semi-automated manner by selecting an intensity threshold above which a spot is considered an RNA particle. The number of puncta, their average size, and total area of signal was calculated for each probe. These values were summed across all images taken from (i) the left and right anterior commissures, (ii) the corpus callosum, or (iii) nine fields of view of the nucleus accumbens (for *Drd1*/IEGs). Area of signal was normalized by cell number, which was quantified by DAPI signal using the NucleusCounter plugin in ImageJ.

### Cnp-Cre/Ribotag dissection and RNA isolation

Cnp-Cre mice were crossed with Rpl22-HA (“RiboTag”)^51^ mice (both in the C57Bl/6 background) to generate Cnp-Cre x RiboTag mice, facilitating the immunoprecipitation of oligodendrocyte-enriched mRNAs for RT-PCR analysis. Mock/morphine treatment and dissection was performed as described above, except that the entire striatum, corpus callosum, or 40 mg of cortex were harvested and transferred to microcentrifuge tubes containing 500 μL polysome lysis buffer. Lysis and immunoprecipitation were performed as previously described^51^. Lysates were applied to Machery Nagel Nucleospin RNA columns and RNA was purified per the manufacturer’s protocol.

### Quantitative real-time PCR

RNA isolated from Cnp-Cre x Ribotag mice or FACS-isolated oligodendrocytes (see below) was reverse transcribed using qScript^™^ cDNA Supermix from QuantaBio according to the manufacturer’s protocol. cDNA was mixed with Fast SYBR^™^ Green master mix from ThermoFisher and primers to the indicated gene (Supplementary Table 8) in a 96-well plate, and quantified using the Quant Studio 3 from Applied Biosciences (in technical triplicates). Differences in relative gene expression were calculated using the delta delta cT method.

### Immunostaining and FACS of oligodendrocytes

WT mice were injected with saline (mock) or morphine as described above. Four h later, mice were euthanized and entire brains were removed, minced with a razor blade, and dissociated with papain solution (400 units of papain in 8 mL total volume). After 50 min dissociation, suspensions were diluted with EBSS #2^44^, serially passed through 100 μm and 40 μm filters, and centrifuged at 300*g* for 5 min. The pellet was resuspended in 35% Percoll solution prepared in EBSS containing 0.1% BSA and 5% trehalose, then centrifuged at 800*g* for 20 min at 18°C. The supernatant containing myelin and cell debris was carefully removed and the pellet was resuspended in flow cytometry buffer (1% normal mouse serum, 1% normal rat serum, 5% trehalose, 0.1% BSA, 1 mM EDTA in DPBS). Cells were counted using a hemocytometer and ~3e6 cells were transferred to a fresh tube, pelleted by brief centrifugation at 300g, and resuspended in 200 μL flow cytometry buffer containing the following fluorophore-conjugated antibodies (or isotype controls): GalC-FITC (4 μg; Millipore), PDGFRa-PE (2 μg; BD Biosciences), Mog-Cy5 (4 μg; Millipore; conjugated with Lightning Link kit from Novus Bio), and CD45-PacBlue (2 μg; BioLegend). Tubes were incubated at 4°C with gentle rotation for 30 min, protected from light. Cells were washed twice with and resuspended in flow cytometry buffer (500 μL), and stored on ice protected from light until sorting (<30 min). Samples were sorted into flow cytometry buffer using the Sony SY3200 “Synergy” with 100 μm nozzle size at the Siteman Cancer Center Flow Cytometry Core.

### RNA purification, library preparation, and sequencing

Immediately following FACS, cells (~100K per sample) were centrifuged at 300*g* for 5 min at 4°C, and the supernatant was carefully removed. The pellet was resuspended in 300 μL Trizol reagent and RNA was purified according to the manufacturer’s protocol. Sequencing libraries were generated using the SeqPlex RNA Amplification Kit (Sigma) according to the manufacturer’s instructions, and sequenced to a depth of ~30 million raw reads per sample (single-end, 50 bp) on the Illumina HiSeq2500.

### RNAseq alignment and DE analysis

Reads were mapped to the mouse genome mm10 using STAR. We used htseq to generate gene-count tables for all samples, then performed differential expression between the 3 replicates each from mock- or morphine-treated mice using DESeq. We first removed genes whose basemean expression was <30, leaving ~12,000 genes. For Fig. 5a, we filtered the genes by adjusted p-value (p-adj) and log_2_(fold-change) to generate two lists with increasing stringency: “Top 1000” - p-adj < 0.05 and abs[log_2_FC] > 0.5; (ii) “Top 200” - p-adj < 1e-12 and abs[log_2_FC] > 1.

### Pathway analysis of single-cell and bulk RNAseq data

Pathway analysis was performed using Genomatix GeneRanker software. For single-cell data, we used as input lists: (i) genes enriched in D1 MSNs of the activated MSN cluster compared to all other D1 MSNs (SCDE p < 0.001; 256 genes), (ii) morphine-regulated genes in oligodendrocytes (MO_all) (SCDE p < 0.05, abs[log_2_FC] > 0.4; 69 genes), or (iii) morphine-regulated genes in astrocytes (SCDE p < 0.3; 81 genes). For bulk RNAseq analysis, we used the top 200 morphine-induced (p-adj < 1e-12, log_2_FC > 1) or the top 148 morphine-repressed (p-adj < 1e-8, log_2_FC < -0.5) genes. The pathway shown in Supplementary Figures 7 and 12 includes genes associated with the pharmacological substance dexamethasone, a corticosteroid and activator of the glucocorticoid receptor (Supplementary Table S7).

## ACKNOWLEDGEMENTS

This work was supported by NIH grants CA009547-33 to DA, AGO13730 and NS099314 to JM, and U54 HD087011 to the Intellectual and Developmental Disabilities Research Center at Washington University. We thank the Alvin J. Siteman Cancer Center at Washington University School of Medicine and Barnes-Jewish Hospital in St. Louis, MO., for the use of the Siteman Flow Cytometry core. The Siteman Cancer Center is supported in part by an NCI Cancer Center Support Grant (#P30 CA091842). We thank Peter Wang and Gwendalyn Randolph for sharing the CD45-PacBlue antibody.

## SUPPLEMENTARY DATA

**Supplementary Figure 1** Quality control of Drop-seq libraries and correlation with previous RNA-seq datasets of the mouse nucleus accumbens.

**Supplementary Figure 2** Proportions and marker genes of major CNS cell types are unchanged by acute morphine treatment.

**Supplementary Figure 3** Markers of neuronal subclusters are consistent with ISH data from the Allen Brain Atlas.

**Supplementary Figure 4** Heatmaps showing single-cell relative expression of neuronal cluster-enriched genes.

**Supplementary Figure 5** Pathway analysis of genes enriched in morphine-treated *Drd1*-expressing cells of the Activated MSN cluster.

**Supplementary Figure 6** Subclustering further resolves non-neuronal cell types of the NAc.

**Supplementary Figure 7** Distinct subsets of glucocorticoid receptor (GR) target genes are induced by morphine in oligodendrocytes (MO_all) and astrocytes (Astrocyte_all).

**Supplementary Figure 8** FISH of *Cdkn1a* in the corpus callosum (outlined by dotted line)

**Supplementary Figure 9** FISH of neuronal genes in the NAc. Arrows in NTX/Morphine Merge panels indicated cells positive for the indicated IEG (*Nr4a1, Junb*, or *Fos*), but negative for *Drd1*.

**Supplementary Figure 10** FACS plots illustrating gating for GALC^+^/MOG^+^ oligodendrocytes.

**Supplementary Figure 11** qRT-PCR analysis of RNAs purified from Cnp-Cre x Ribotag mice (**a**) or from FACS-isolated OLs of WT mice (**b**).

**Supplementary Figure 12** Pathway analysis of morphine-induced genes in oligodendrocytes as assessed by bulk RNAseq.

**Supplementary Table 1** Numbers and ratios of cells in each all-cell t-SNE cluster.

**Supplementary Table 2** Top 50 genes enriched in each all-cell t-SNE cluster sorted by FDR. The top 5 enriched genes per cluster are bolded, and are the ones included in the heatmaps in Fig 2.

**Supplementary Table 3** Numbers and ratios of cells in each neuronal t-SNE subcluster.

**Supplementary Table 4** Top 50 genes enriched in each t-SNE subcluster sorted by FDR.

**Supplementary Table 5** Numbers and ratios of cells in each non-neuronal t-SNE subcluster.

**Supplementary Table 6** SCDE result of every cell type cluster.

**Supplementary Table 7** Pathway analysis results of morphine-regulated genes in activated MSNs, oligodendrocytes, and astrocytes identified by Drop-seq/SCDE.

**Supplementary Table 8** List of primers used for qRT-PCR.

**Supplementary Table 9** Full list of genes significantly affected by morphine in oligodendrocytes as assessed by bulk RNA-seq.

**Supplementary Table 10** Pathway analysis results of morphine-regulated genes in oligodendrocytes identified by FACS-RNA-seq.

## REFERENCES

1. Bauer, I.E., Soares, J.C. & Nielsen, D.A. The role of opioidergic genes in the treatment outcome of drug addiction pharmacotherapy: A systematic review. Am JAddict 24, 15–23 (2015).

2. Dart, R.C., et al. Trends in opioid analgesic abuse and mortality in the United States. NEngl J Med 372, 241–248 (2015).

3. Liu, H., et al. Disrupted white matter integrity in heroin dependence: a controlled study utilizing diffusion tensor imaging. Am J Drug Alcohol Abuse 34, 562–575 (2008).

4. Bora, E., et al. White matter microstructure in opiate addiction. Addict Biol 17, 141–148 (2012).

5. Luscher C. & Malenka R.C. Drug-evoked synaptic plasticity in addiction: from molecular changes to circuit remodeling. Neuron 69, 650–663 (2011).

6. Volkow N.D. & Morales M. The Brain on Drugs: From Reward to Addiction. Cell 162, 712–725 (2015).

7. Zhu, L., et al. Chronic methamphetamine regulates the expression of MicroRNAs and putative target genes in the nucleus accumbens of mice. JNeurosci Res 93, 1600–1610 (2015).

8. Piechota, M., et al. The dissection of transcriptional modules regulated by various drugs of abuse in the mouse striatum. Genome Biol 11, R48 (2010).

9. Albertson, D.N., Schmidt, C.J., Kapatos, G. & Bannon, M.J. Distinctive profiles of gene expression in the human nucleus accumbens associated with cocaine and heroin abuse. Neuropsychopharmacology 31, 2304–2312 (2006).

10. Korostynski, M., Piechota, M., Kaminska, D., Solecki, W. & Przewlocki, R. Morphine effects on striatal transcriptome in mice. Genome Biol 8, R128 (2007).

11. Islam, S., et al. Quantitative single-cell RNA-seq with unique molecular identifiers. Nat Methods 11. 163–166 (2014).

12. Cao, J., et al. Comprehensive single-cell transcriptional profiling of a multicellular organism. Science 357, 661–667 (2017).

13. Klein, A.M., et al. Droplet barcoding for single-cell transcriptomics applied to embryonic stem cells. Cell 161, 1187–1201 (2015).

14. Zheng, G.X., et al. Massively parallel digital transcriptional profiling of single cells. Nat Commun 8, 14049 (2017).

15. Macosko, E.Z., et al. Highly Parallel Genome-wide Expression Profiling of Individual Cells Using Nanoliter Droplets. Cell 161, 1202–1214 (2015).

16. van der Maaten, L.H., G. Visualizing data using t-SNE. J. Mach. Learn. Res. 9, 2579–2605 (2008).

17. Gokce, O., et al. Cellular Taxonomy of the Mouse Striatum as Revealed by Single-Cell RNA-Seq. Cell Rep 16, 1126–1137 (2016).

18. Zhang, Y., et al. An RNA-sequencing transcriptome and splicing database of glia, neurons, and vascular cells of the cerebral cortex. J Neurosci 34, 11929–11947 (2014).

19. Hodes, G.E., et al. Sex Differences in Nucleus Accumbens Transcriptome Profiles Associated with Susceptibility versus Resilience to Subchronic Variable Stress. JNeurosci 35, 16362–16376 (2015).

20. Lobo, M.K., Karsten, S.L., Gray, M., Geschwind, D.H. & Yang, X.W. FACS-array profiling of striatal projection neuron subtypes in juvenile and adult mouse brains. Nat Neurosci 9, 443–452 (2006).

21. Enoksson, T., Bertran-Gonzalez, J. & Christie, M.J. Nucleus accumbens D2- and D1-receptor expressing medium spiny neurons are selectively activated by morphine withdrawal and acute morphine, respectively. Neuropharmacology 62, 2463–2471 (2012).

22. Marques, S., et al. Oligodendrocyte heterogeneity in the mouse juvenile and adult central nervous system. Science 352, 1326–1329 (2016).

23. Kharchenko, P.V., Silberstein, L. & Scadden, D.T. Bayesian approach to single-cell differential expression analysis. Nat Methods 11, 740–742 (2014).

24. Marinelli, M., Aouizerate, B., Barrot, M., Le Moal, M. & Piazza, P.V. Dopamine-dependent responses to morphine depend on glucocorticoid receptors. Proc Natl Acad Sci U S A 95, 7742–7747 (1998).

25. Slezak, M., et al. Astrocytes are a neural target of morphine action via glucocorticoid receptor-dependent signaling. Glia 61, 623–635 (2013).

26. Dal Molin, A., Baruzzo, G. & Di Camillo, B. Single-Cell RNA-Sequencing: Assessment of Differential Expression Analysis Methods. Front Genet 8, 62 (2017).

27. Robinson, A.P., Rodgers, J.M., Goings, G.E. & Miller, S.D. Characterization of oligodendroglial populations in mouse demyelinating disease using flow cytometry: clues for MS pathogenesis. PLoS One 9, e107649 (2014).

28. Lin W. & Popko B. Endoplasmic reticulum stress in disorders of myelinating cells. Nat Neurosci 12, 379–385 (2009).

29. Upadhyay, J., et al. Alterations in brain structure and functional connectivity in prescription opioid-dependent patients. Brain 133, 2098–2114 (2010).

30. Wang, Y., et al. White matter impairment in heroin addicts undergoing methadone maintenance treatment and prolonged abstinence: a preliminary DTI study. Neurosci Lett 494, 49–53 (2011).

31. Li, W, et al. Brain white matter integrity in heroin addicts during methadone maintenance treatment is related to relapse propensity. Brain Behav 6, e00436 (2016).

32. Knapp, P.E., Maderspach, K. & Hauser, K.F. Endogenous opioid system in developing normal and jimpy oligodendrocytes: mu and kappa opioid receptors mediate differential mitogenic and growth responses. Glia 22, 189–201 (1998).

33. Stiene-Martin, A., et al. Opioid system diversity in developing neurons, astroglia, and oligodendroglia in the subventricular zone and striatum: impact on gliogenesis in vivo. Glia 36, 78–88 (2001).

34. Sanchez, E.S., Bigbee, J.W., Fobbs, W., Robinson, S.E. & Sato-Bigbee, C. Opioid addiction and pregnancy: perinatal exposure to buprenorphine affects myelination in the developing brain. Glia 56, 1017–1027 (2008).

35. Vestal-Laborde, A.A., Eschenroeder, A.C., Bigbee, J.W., Robinson, S.E. & Sato-Bigbee, C. The opioid system and brain development: effects of methadone on the oligodendrocyte lineage and the early stages of myelination. Dev Neurosci 36, 409–421 (2014).

36. Simon, M., George, R. & Garcia, J. Acute morphine effects on regional brain amines, growth hormone and corticosterone. Eur J Pharmacol 34, 21–26 (1975).

37. Miyata, S., et al. Plasma corticosterone activates SGK1 and induces morphological changes in oligodendrocytes in corpus callosum. PLoS One 6, e19859 (2011).

38. Miyata, S., Hattori, T., Shimizu, S., Ito, A. & Tohyama, M. Disturbance of oligodendrocyte function plays a key role in the pathogenesis of schizophrenia and major depressive disorder. Biomed Res Int 2015, 492367 (2015).

39. Ammon-Treiber, S., et al. Rapid, transient, and dose-dependent expression of hsp70 messenger RNA in the rat brain after morphine treatment. Cell Stress Chaperones 9, 182–197 (2004).

40. Salas, E., et al. Gene expression analysis of heat shock proteins in the nucleus accumbens of rats with different morphine seeking behaviours. Behav Brain Res 225, 71–76 (2011).

41. Volpi, V.G., Touvier, T. & D’Antonio, M. Endoplasmic Reticulum Protein Quality Control Failure in Myelin Disorders. FrontMolNeurosci 9, 162 (2016).

42. Hussien, Y., et al. ER Chaperone BiP/GRP78 Is Required for Myelinating Cell Survival and Provides Protection during Experimental Autoimmune Encephalomyelitis. JNeurosci 35, 15921–15933 (2015).

43. Eschenroeder, A.C., Vestal-Laborde, A.A., Sanchez, E.S., Robinson, S.E. & Sato-Bigbee, C. Oligodendrocyte responses to buprenorphine uncover novel and opposing roles of mu-opioid- and nociceptin/orphanin FQ receptors in cell development: implications for drug addiction treatment during pregnancy. Glia 60, 125–136 (2012).

44. Saxena, A., et al. Trehalose-enhanced isolation of neuronal sub-types from adult mouse brain. Biotechniques 52, 381–385 (2012).

45. Campbell, J.N., et al. A molecular census of arcuate hypothalamus and median eminence cell types. Nat Neurosci 20, 484–496 (2017).

46. McCarthy, D.J., Campbell, K.R., Lun, A.T. & Wills, Q.F. Scater: pre-processing, quality control, normalization and visualization of single-cell RNA-seq data in R. Bioinformatics 33, 1179–1186 (2017).

47. At, L.L., Bach, K. & Marioni, J.C. Pooling across cells to normalize single-cell RNA sequencing data with many zero counts. Genome Biol 17, 75 (2016).

48. Satija, R., Farrell, J.A., Gennert, D., Schier, A.F. & Regev, A. Spatial reconstruction of singlecell gene expression data. Nat Biotechnol 33, 495–502 (2015).

49. Butler, A., Hoffman, P., Smibert, P., Papalexi, E. & Satija, R. Integrating single-cell transcriptomic data across different conditions, technologies, and species. Nat Biotechnol (2018).

50. Schindelin, J., et al. Fiji: an open-source platform for biological-image analysis. Nat Methods 9, 676–682 (2012).

51. Sanz, E., et al. Cell-type-specific isolation of ribosome-associated mRNA from complex tissues. Proc Natl Acad Sci U S A 106, 13939–13944 (2009).

